# Niche cells regulate primordial germ cell quiescence in response to basement membrane signaling

**DOI:** 10.1101/2023.02.05.527217

**Authors:** Daniel C. McIntyre, Jeremy Nance

**Affiliations:** Skirball Institute of Biomolecular Medicine, NYU Grossman School of Medicine, New York, NY 10016; Department of Cell Biology, NYU Grossman School of Medicine, New York, NY 10016; University of Virginia, Department of Biology, 90 Geldard Drive, Physical Life Science Building Room 318, Charlottesville, VA 22904

**Keywords:** primordial germ cell, niche, perlecan, collagen, integrin, laminin

## Abstract

Stem cell quiescence, proliferation and differentiation are controlled by interactions with niche cells and a specialized extracellular matrix called the basement membrane (BM). Direct interactions with adjacent BM are known to regulate stem cell quiescence; however, it is less clear how niche BM relays signals to stem cells that it does not contact. Here, we examine how niche BM regulates *C. elegans* primordial germ cells (PGCs), which remain quiescent during embryogenesis. Depleting the BM protein laminin causes embryonic PGCs to proliferate, indicating that laminin functions to maintain PGC quiescence. How laminin signals to the PGCs remains unclear, as somatic niche cells enwrap PGCs and physically exclude them from contacting the BM. Here, we show that, following laminin depletion, gonadal niche cells relay proliferation-inducing signals from the gonadal BM to PGCs via integrin receptors. Mutations disrupting the BM proteoglycan perlecan block PGC proliferation when laminin is depleted, suggesting that laminin functions to inhibit a proliferation-inducing signal originating from perlecan. Our results reveal how BM signals can regulate stem cell quiescence indirectly, by activating niche cell integrin receptors.

## Introduction

Stem cells must balance proliferation and differentiation in order to meet the developmental and regenerative needs of an organism. To accomplish such precise regulation, stem cells rely on information conveyed from a surrounding micro- environment called the niche. The niche is composed of both cells and extracellular matrix (ECM) (Scadden, 2014, Jones and Wagers, 2008). Niche cells and ECM can function by interacting directly with stem cells. In addition, ECM can impact stem cells indirectly, either by acting through niche cells or by modulating the local signaling environment. Indirect control of stem cells by niche ECM adds an additional, underappreciated layer of complexity to niche function.

Basement membrane (BM) is one common type of extracellular matrix contributing to stem cell niches (Kruegel and Miosge, 2010). For example, mammalian muscle, epidermal, neuronal, hematopoietic and intestinal stem cells are all closely juxtaposed to BMs (Gattazzo et al., 2014). BMs are thin flexible sheets of extracellular matrix that include four core components conserved in all bilaterian animals: sheet-like layers of laminin and collagen IV; the proteoglycan perlecan; and the crosslinking protein nidogen (Yurchenco, 2011, Pastor-Pareja, 2020). These four components form a matrix (the LNCP matrix) that cells sense using cell-surface receptors including integrin, dystroglycan and glypican (Yurchenco, 2011). Numerous additional proteins associate with the core BM matrix, altering its composition in different regions of the organism and during various developmental stages (Yurchenco, 2011, Jayadev et al., 2019).

Although BMs function as physical barriers that support and separate adjacent tissues, they also provide critical signaling inputs to surrounding cells. Core BM proteins can signal to adjacent cells by interacting with cell surface BM receptors. For example, collagen IV and perlecan contain Arg-Gly-Asp (RGD) motifs than can activate a specific class of RGD-binding integrin receptors (Ludwig et al., 2021). Other types of integrins bind the laminin G-like (LG) domains present in laminin α subunits (Yurchenco, 2011). Such interactions can relay both biochemical and mechanical properties of the BM to cells. In addition, BM proteins can affect local signaling indirectly by binding to a wide variety of growth factors. For example, in addition to binding to integrin directly through its RGD motifs, perlecan binds directly to FGF, Wnt and TGF-ß ligands, affecting output from these signaling pathways in nearby cells by either concentrating or sequestering ligands (Hayes et al., 2022). It is clear from a variety of model systems that the signaling capacity of BMs is essential for stem cell niche function. For example, BM signaling ensures self-renewal of embryonic stem cells and hair follicle stem cells in mammals, and follicular stem cells in the fly ovary (Suh and Han, 2011, O’Reilly et al., 2008, Fujiwara et al., 2011). Since integrin signaling also controls proliferation in many types of cancer (Cooper and Giancotti, 2019), one possibility is that dysfunctional tumor BM could employ similar signaling mechanisms to regulate cancer stem cells.

One of the most important niche functions is balancing stem cell quiescence and proliferation. Failure to properly regulate proliferation can lead to developmental defects and contributes to aging and cancer growth (Oh et al., 2014, Kai et al., 2019). This regulation often occurs through cooperation between niche BM and growth factor signaling ligands. For example, proliferation of mouse neural stem cells is controlled by perlecan, which binds growth factors such as FGF2 (Sirko et al., 2007, Kerever et al., 2007, Kerever et al., 2014). Alternatively, transduction of BM signals by integrin receptors can impact stem cell proliferation. For example, in mouse muscle stem cells β1 integrin potentiates FGF2 signaling, which drives expansion of muscle stem cells (Rozo et al., 2016). Indeed, β1 integrin regulates proliferation across multiple mammalian tissues including the skin, mammary and intestinal epithelium as well as neural stem cells in the subventricular zone (Shen et al., 2008, Raghavan et al., 2000, Jones et al., 2006, Faraldo et al., 2001). Integrin receptors also regulate stem cell proliferation in flies, often by cooperating with other signaling pathways, as reported for intestinal stem cells, follicular stem cells and blood progenitors (O’Reilly et al., 2008, Lin et al., 2013, Khadilkar et al., 2020). Thus, niche BM, and integrin signal transduction in particular, are critical regulators of stem cell proliferation.

In principle, information from the BM can influence stem cell behavior in two ways. First, signals may flow directly from the BM to stem cells, either through BM receptors on stem cells contacting the matrix or through modulation of secreted signals by the BM. Alternatively, niche cells can interact with the BM and relay information to stem cells. In both the *C. elegans* and *D. melanogaster* germ lines, niche cells act as intermediaries between stem cells and BM. In fly ovaries, formation of the gonadal BM is needed for normal niche cell morphology and for BMP signaling between niche and stem cells. Disrupting collagen IV deposition results in ovaries with too many germline stem cells (Van De Bor et al., 2015, Diaz-Torres et al., 2021). Similarly, in the *C. elegans* embryonic gonad primordium, we showed previously that depleting laminin severely disrupts niche morphology and induces germ cell proliferation (McIntyre and Nance, 2020). The heparan sulfate proteoglycan syndecan also functions in *C. elegans* gonadal niche cells to modulate germline stem cell proliferation (Gopal et al., 2021). However, in these systems the mechanisms niche cells use to convey information from the BM to stem cells remains largely unknown.

In this study, we investigate how niche BM and niche cells in the *C. elegans* gonad primordium collaborate to control quiescence of primordial germ cells (PGCs), which are the direct precursors of germline stem cells. The gonad primordium assembles during embryogenesis when two somatic gonadal precursor cells (SGPs), which function as niche cells, enwrap the two PGCs to form a four-cell gonad primordium (Fig. 1A) (Rohrschneider and Nance, 2013, Sulston et al., 1983). Soon after the gonad primordium assembles, SGPs nucleate a BM that surrounds the gonad primordium (Sibley et al., 1993, McIntyre and Nance, 2020, Kramer, 2000, Kao et al., 2006, Huang et al., 2003, Guo et al., 1991, Graham et al., 1997). Embryonic PGCs are quiescent (arrested in G2 phase) and do not reenter the cell cycle unless the hatched larva encounters food (Fukuyama et al., 2006). We showed previously that depleting laminin from the gonadal BM activates Notch signaling in PGCs, causing them to exit from quiescence precociously and proliferate in the embryo (McIntyre and Nance, 2020). However, it remained unclear from this study how BM signaled to the PGCs. Here, we identify BM proteins that are required for regulating PGC quiescence and show that these signals are relayed to PGCs by gonadal niche cells using Arg-Gly-Asp (RGD) binding integrin receptors. Our data support a model wherein niche cells induce stem cells to exit from quiescence in response to signals from the gonadal BM.

**Figure 1.**
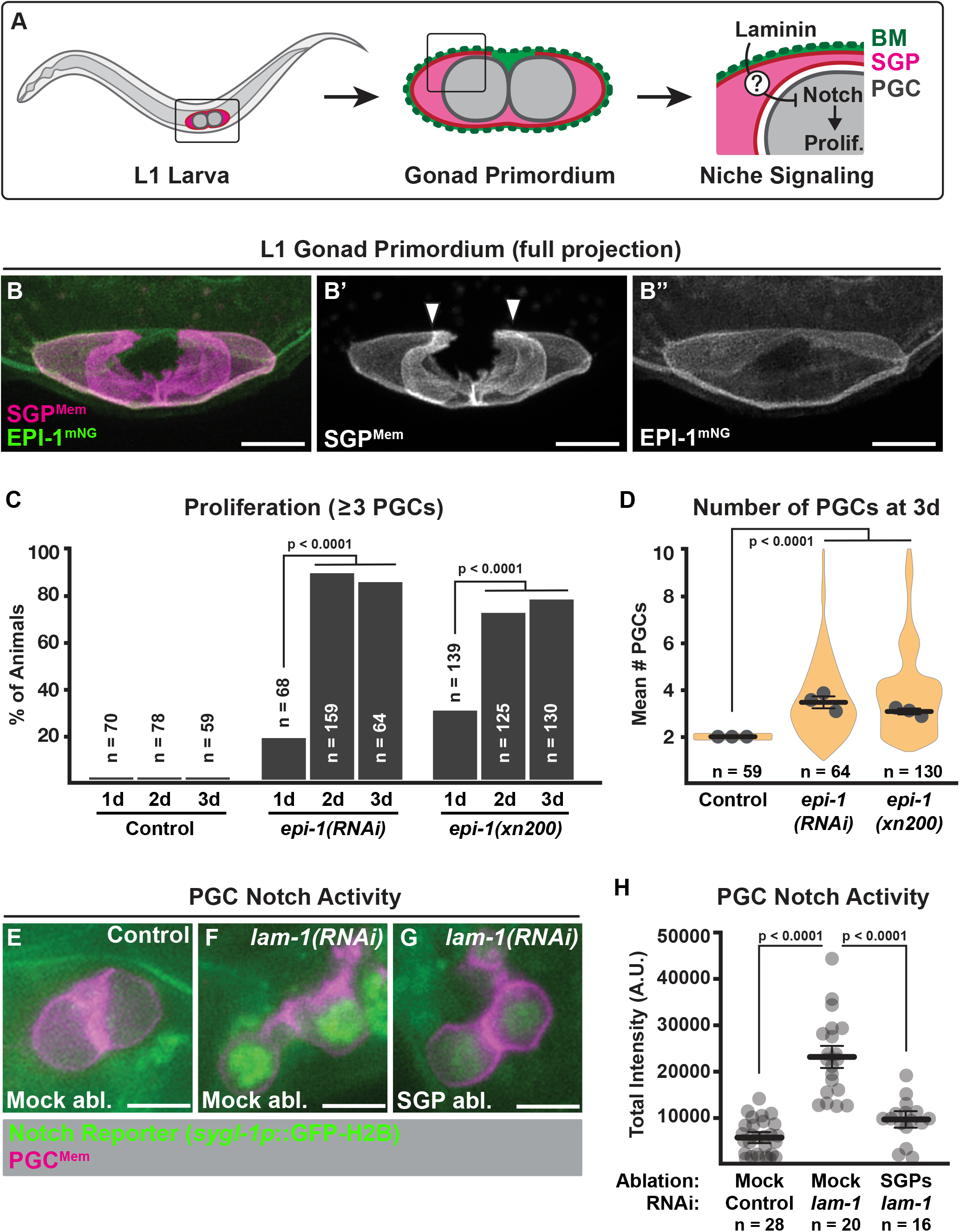
Niche cells transduce basement membrane signals that regulate PGC quiescence. **A**. Basement membrane (BM, green) surrounds the gonad primordium in L1 stage larvae. Laminin in the BM regulates when PGCs exit quiescence and begin to proliferate by inhibiting Notch signaling. Somatic gonadal cells (SGPs, magenta) are positioned to transmit BM signals to PGCs. **B**. The L1 gonad primordium including SGP cell membranes (magenta) and laminin α/EPI-1 (green). Arrowheads indicate the approximate position of the PGCs. **C**. Quantification of the fraction of starved L1 larvae with proliferating PGCs for each genotype and time in days (d) post-hatching. **D**. Average number of PGCs for each for each genotype in starved L1 larvae. **E – G**. Notch transcriptional reporter activity (green) in PGCs (magenta). Genotype and SGP ablation are as indicated. **H**. Quantification of total nuclear GFP fluorescence intensity in PGCs from E – G. Scale bars are 5μm and specific p-values are listed (n.s., not significantly different, p > 0.05). Fisher’s exact test was used to compare the faction of worms with proliferating PGCs in C. All other quantitative comparisons used a 2-tailed t-test.

## Results

### Laminin depletion induces continued PGC proliferation

Laminin, which is a heterotrimer composed of α, ß and γ subunits, can be removed by inhibiting the sole laminin ß-encoding gene, *lam-1* (Kramer, 2000). Eliminating laminin using a *lam-1* null mutation causes only a fraction of PGCs to exit quiescence, and those that do typically divide only once (McIntyre and Nance, 2020). One potential explanation for this finding is that the gonadal BM regulates only the initial escape from PGC quiescence, after which point another pathway controls continued proliferation. Alternatively, since *lam-1* embryos arrest with major morphogenetic defects (Kramer, 2000), it is possible that the attendant deterioration of mutant embryos prevents additional rounds of PGC proliferation. To distinguish between these possibilities, we inhibited the laminin α gene *epi-1*, which encodes the primary laminin α detected in the gonadal BM (Fig. 1B) (McIntyre and Nance, 2020). *epi-1* functions redundantly with the only other laminin α gene *(lam-3)* in embryonic development; consequently, in contrast to *lam-1* mutant embryos, most *epi-1(RNAi)* embryos are able to hatch (95/103 embryos). *epi-1(RNAi)* L1 larva starved for two or three days showed significantly greater PGC proliferation than those starved only a single day (Fig. 1C), and ∼20% of larvae starved for three days contained more than four PGCs (up to 10 PGCs, Fig. 1D). We observed comparable results using the *epi-1(xn200)* putative null mutation, which we created by inserting a stop cassette into the second exon of an endogenously tagged *epi-1::mNG* allele (Fig. S1A). Almost all *epi-1(xn200)* mutants died as embryos or young larva (Fig. S1D-F), and no EPI-1^mNG^ was visible in mutant animals (Fig. S1B,C). PGCs in starved *epi- 1(xn200)* L1 larvae underwent a similar number of divisions as those in starved *epi- 1(RNAi)* L1 larvae (Fig. 1C,D). We conclude that depleting laminin from the gonadal BM induces PGCs to exit from quiescence and to continue dividing, perhaps until they can no longer do so because of nutrient limitation.

### Niche cells relay quiescence-regulating signals from the BM to the PGCs

SGP cell membranes enwrap the PGCs, making it unlikely that PGCs make significant contact with the gonadal BM (Rohrschneider and Nance, 2013, McIntyre and Nance, 2020). In both confocal reconstructions of the L1 gonad (Fig. 1B) and published transmission electron micrographs (Sulston et al., 1983), PGCs are separated from the BM by niche cell membranes. Therefore, we hypothesized that SGPs relay laminin-dependent quiescence-regulating signals from the basement membrane to the PGCs. We tested this hypothesis using laser microsurgery to kill the SGP ancestors combined with a transgene, *sygl-1p::GFP-H2B*, that reports on Notch signaling activity in the PGCs (Kershner et al., 2014); because *sygl-1p::GFP-H2B* is activated in PGCs when laminin is depleted, and *glp-1/Notch* is required for PGC proliferation under these conditions, *sygl- 1p::GFP-H2B* expression in PGCs is an indicator of exit from quiescence (McIntyre and Nance, 2020). In contrast to mock-ablated *lam-1(RNAi)* embryos, which showed robust activation of *sygl-1p::GFP-H2B*, PGCs in ablated *lam-1(RNAi)* embryos lacking SGPs failed to activate *sygl-1p::GFP-H2B* (Fig. 1E–H). These results demonstrate that SGPs are required to activate Notch signaling in PGCs when laminin is depleted, and are consistent with classic laser ablation experiments showing that SGPs are needed for PGCs to proliferate in fed larva (Kimble and White, 1981). Based on these findings, we propose that SGPs relay signals from the BM to the PGCs, regulating their exit from quiescence (Fig. 1A).

### Niche cell integrins transduce quiescence-regulating BM signals

We hypothesized that SGPs express a receptor that transduces BM signals. Integrin BM receptors function as heterodimers comprised of α and ß subunits. *C. elegans* has two integrin α genes (*pat-2* and *ina-1*) and a single integrin ß gene (*pat-*3) (Williams and Waterston, 1994, Gettner et al., 1995, Baum and Garriga, 1997). SGPs profiled by single-cell RNA sequencing contain transcripts for all three genes (Packer et al., 2019), indicating that SGPs could express α/ß integrins INA-1/PAT-3 and PAT- 2/PAT-3 at their surfaces. To assess integrin protein localization and function in the SGPs, we tagged the endogenous *pat-3* gene with sequences encoding GFP and the ZF1 degron, which can be used to conditionally deplete proteins (*xn109[pat-3::gfp::zf1]*; Fig. S2A) (Armenti et al., 2014). In addition to robust expression in other tissues contacting BM, including the pharyngeal primordium and body-wall muscles, PAT-3^GFP- ZF1^ was present on SGP cell membranes in both embryonic and L1 gonad primordia (Fig. 2A,B). PAT-3^GFP-ZF1^ remained on SGP cell surfaces in *epi-1(RNAi)* embryos, although at slightly reduced levels (Fig. 2C–E), indicating that integrin is appropriately positioned to transduce signals from the BM to SGPs following laminin depletion.

**Figure 2.**
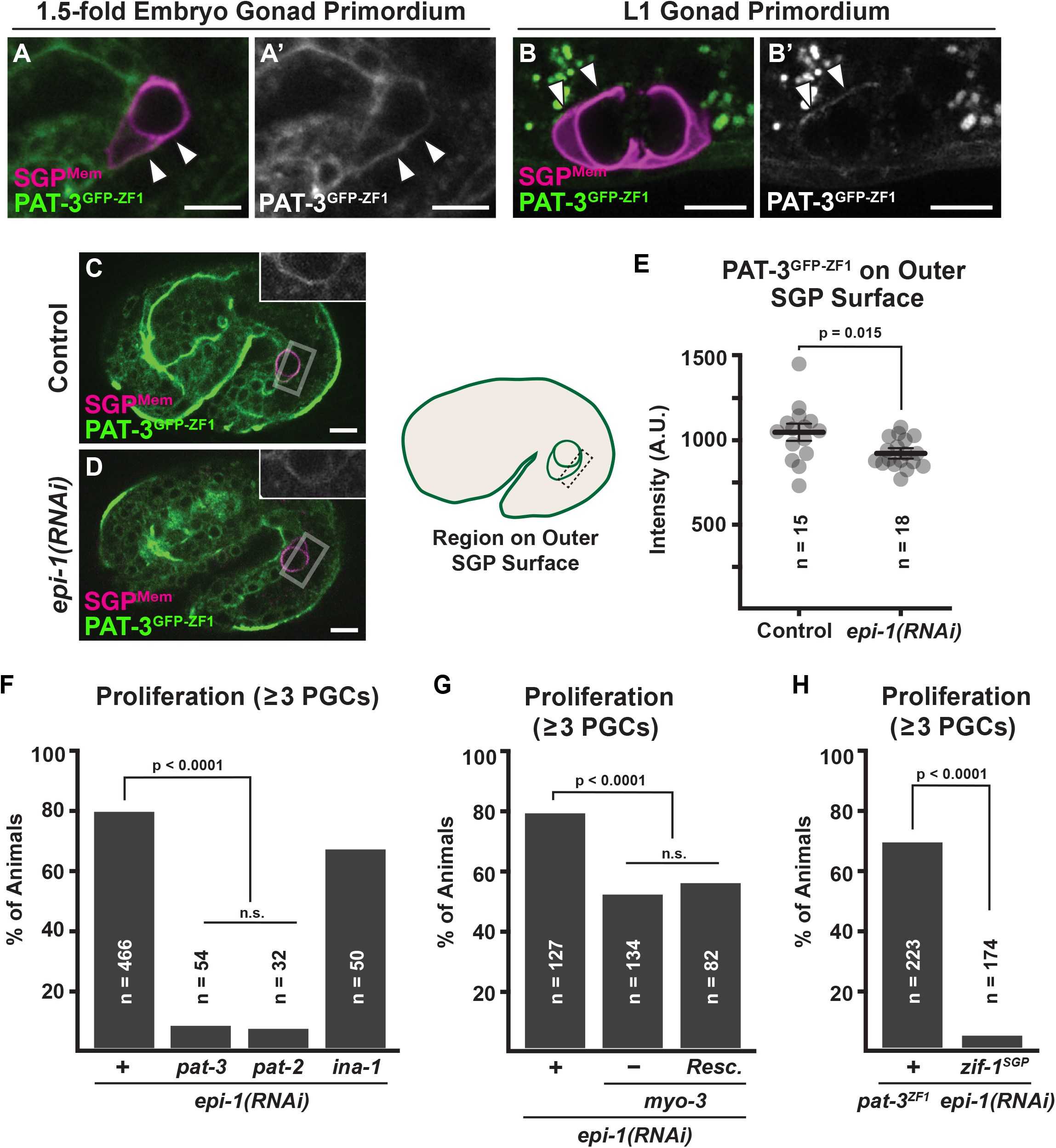
Integrin within SGPs is necessary for PGCs to exit quiescence when laminin is depleted. **A – B**. Integrin ß/PAT-3^GFP-ZF1^ (green) expression on outer surfaces of SGPs (magenta) during gonad development (A) and in L1 larvae (B). **C – E**. Integrin ß/PAT-3^GFP-ZF1^ (green) expression in control (C) and *epi-1(RNAi)* (D) embryos. Integrin ß/PAT-3^GFP-ZF1^ expression on SGP membranes (boxed insets in C and D) is quantified in E. **F – H**. Quantification of the fraction of starved L1 larvae with proliferating PGCs for the genotypes indicated (*myo-3* rescued by a *myo-3(+)* extrachromosomal array). Scale bars are 5μm and specific p-values are listed (n.s., not significantly different, p > 0.05). Fisher’s exact test was used to compare the fraction of worms with proliferating PGCs in F – H. All other quantitative comparisons used a 2-tailed t-test.

To examine integrin function in PGC quiescence, we engineered a putative null allele of *pat-3* by inserting a stop cassette into the *pat-3(xn109[pat-3::gfp::zf1])* allele upstream of the sequence encoding the transmembrane domain [*pat-3(xn113)*, Fig. S2B]. Like existing strong *pat-3* mutations (Williams and Waterston, 1994), *pat-3(xn113)* embryos arrested at the two-fold stage of embryogenesis (54/55 embryos). We first assessed whether *pat-3* mutant embryos, like those depleted of laminin, also showed inappropriate PGC proliferation. Contrary to our expectation if laminin signaled through integrin receptors, PGCs did not proliferate in starved *pat-3* mutants (n = 83/83 ≤2 PGCs). Existing alleles of integrin α genes *ina-1* and *pat-2* that are likely null mutations also caused no PGC proliferation phenotype (*ina-1(gm86)* n = 59/59 ≤2 PGCs; *pat- 2(ok2148)* n = 33/33 ≤2 PGCs). These experiments indicate that laminin does not inhibit PGC proliferation by engaging integrin receptors on SGPs.

We next considered an alternative hypothesis: laminin could regulate PGC quiescence by *preventing* activation of integrin receptors, for example by sequestering or inhibiting an integrin ligand. In this case, loss of integrin should suppress the PGC proliferation that occurs when laminin α/*epi-1* is depleted. Consistent with this model, we found that most PGCs did not proliferate in *pat-3(xn113) epi-1(RNAi)* mutants (Fig. 2F). One possible confounding explanation for these results is if the two-fold arrest of *pat-3* mutant embryos is not conducive for PGC proliferation. To address this possibility, we examined *myo-3(st386)* mutants, which lack functional muscle myosin and also arrest at the two-fold stage (Williams and Waterston, 1994). *myo-3(st386)* mutants only modestly suppressed PGC proliferation in *epi-1(RNAi)* embryos (Fig. 2G). Because the mild suppression was equivalent in *myo-3(st386)* mutants that did not arrest because they carried a *myo-3(+)* extrachromosomal array (Fig. 2G), this small effect is likely explained by genetic background and is not a result of failure in elongation. These findings show that integrin receptors regulate PGC proliferation.

To determine which integrin heterodimer transduces the BM signal, we depleted laminin α/*epi-1* in *ina-1* and *pat-2* mutants. Whereas PGC proliferation in *ina-1 epi- 1(RNAi)* mutant larvae was similar to that in *epi-1(RNAi)* larvae, almost no PGC proliferation occurred in *pat-2 epi-1(RNAi)* mutants (Fig. 2F). Together, these findings suggest that EPI-1 laminin trimers in the BM normally inhibit PGC proliferation by preventing another BM component from activating PAT-2/PAT-3 integrin receptors. This model also agrees with the ligand specificities for the two α integrin subunits in *C. elegans*: INA-1/PAT-3 dimers are predicted to bind to laminin, which is depleted in these experiments, whereas PAT-2/PAT-3 dimers are predicted to bind Arg-Gly-Asp (RGD) sequences present in other BM proteins such as collagen IV and perlecan (Williams and Waterston, 1994, Jayadev et al., 2019, Gettner et al., 1995).

We tested whether integrin function is required in SGPs by utilizing the ZF1 degron we engineered into the *pat-3(xn109[pat-3::gfp::zf1])* allele. Proteins tagged with the ZF1 domain can be rapidly degraded by conditionally expressing the ZIF-1 E3 ligase substrate adaptor (Armenti et al., 2014, Abrams and Nance, 2021). To achieve SGP-specific degradation of PAT-3^GFP-ZF1^, we expressed ZIF-1 from the *ehn-3* promoter (ZIF- 1^SGP^), which is active only in SGPs from the time of gonad formation to late embryogenesis (Mathies et al., 2003). We tested the specificity of ZIF-1^SGP^ by examining ubiquitously expressed transgenic CDC-42^GFP-ZF1^, which was specifically degraded in SGPs (Fig. S2C). Alone, *pat-3*^*GFP-ZF1*^ did not suppress PGC proliferation induced by *epi- 1* RNAi (Fig. 2H). However, when *pat-3*^*GFP-ZF1*^ was combined with ZIF-1^SGP^ in *epi-1(RNAi)* animals, PGC proliferation was suppressed to a similar extent as in *epi-1(RNAi) pat- 3(xn113)* null mutants (Fig. 2H). Because *pat-3*^*GFP-ZF1*^; *zif-1*^*SGP*^ animals do not arrest during embryogenesis like *pat-3* and *pat-2* mutants, this experiment further demonstrates that the suppression of PGC proliferation in *epi-1(RNAi)* larvae is not an indirect consequence of embryonic arrest, but rather results from specific loss of integrin function in the SGPs. All together, these findings indicate that SGP integrins are required to transduce quiescence-regulating signals from the BM to the PGCs.

### Core components of the gonadal BM assemble sequentially

The experiments above raise the possibility that a BM protein other than laminin activates PAT-2/3 integrin receptors on SGPs when laminin α/*epi-1* is depleted. To determine if any of the other three core BM proteins in the LNCP matrix (nidogen, collagen IV, and perlecan) could function as an integrin ligand to relay quiescence signals to the PGCs, we first examined their expression in the gonadal BM. Using a recently described collection of BM genes endogenously tagged to produce mNeonGreen (mNG) fusion proteins (Keeley et al., 2020), we imaged tagged laminin α (EPI-1^mNG^), the sole nidogen (NID-1^mNG^), one of the two subunits of collagen IV (α1 EMB-9^mNG^; the α2 subunit is LET-2) and the sole perlecan (UNC-52^mNG^) in live embryos during gonad formation. Because each of these proteins is fused with the same fluorescent protein, differences in their order of appearance likely reflect real properties of the gonadal BM. LNCP matrix proteins appeared to be added to the gonadal BM in three waves. Laminin and nidogen assembled first, surrounding the outer surfaces of the SGPs once they began wrapping the PGCs (Fig. 3B–E, F–I; yellow rectangles highlight gonadal BM accumulation). Collagen IV became visible in the gonadal BM ∼60 min. after laminin and nidogen first appeared (Fig. 3J – 3M; yellow rectangles highlight accumulation beginning at the 1.5- fold stage). Finally, perlecan accumulated approximately 30 minutes later (Fig. 3N–Q, yellow rectangles highlight accumulation beginning at the 2-fold stage). This ordered accumulation (L + N > C > P; Fig. 3A) mirrors previous findings in other *C. elegans* basement membranes (Keeley et al., 2020), and demonstrates that all parts of the LNCP matrix are present in the gonadal BM by the 2-fold stage and thus could contribute to PGC proliferation.

**Figure 3.**
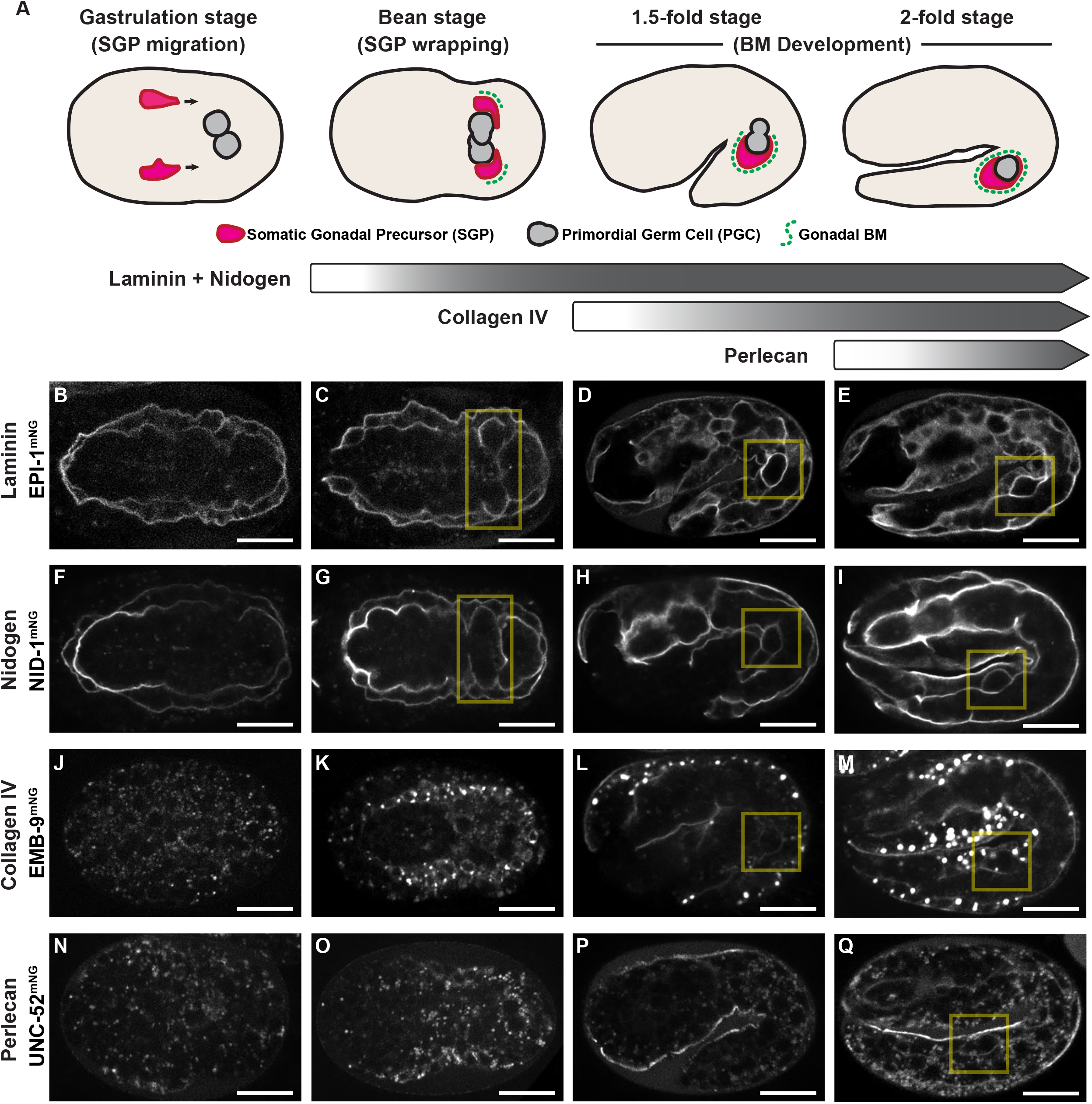
Core components of the gonadal BM assemble sequentially. **A**. Timing of gonadal BM assembly during embryo development from gastrulation stage through the, 2-fold stage. Shaded bars indicate the timing of each core BM component’s incorporation into the gonadal BM. **B – Q**. Expression of core BM components at gastruation, bean, 1.5-fold and 2-fold stages. Images correspond with embryos diagrammed above in A. Yellow boxes highlight expression in the gonadal BM. B – E, laminin /EPI-1^mNG^. F – I, nidogen/NID-1^mNG^. J – M, collagen IV 1/EMB-9^mNG^. N – Q, perlecan/UNC-52^mNG^. Scale bars are 5μm.

### A perlecan RGD motif regulates PGC proliferation

We hypothesized that another BM component activates integrin receptors in SGPs when laminin α/*epi-1* is depleted. As a first test to determine which BM components could activate integrin in the SGPs following laminin knock down, we assayed whether type IV collagen, nidogen and perlecan were still present in the gonadal BM in *epi-1(RNAi)* L1 larvae. First, we examined collagen IV α1/EMB-9^mNG^ localization. In newly hatched, starved L1 larvae, EMB-9^mNG^ was present in the gonadal BM of both control and *epi-1(RNAi)* larva (Fig. 4A,B). Although EMB-9^mNG^ localization in *epi-1(RNAi)* L1 larva was discontinuous, quantification of fluorescence intensity showed no significant change in the amount of EMB-9^mNG^ present in the gonadal BM adjacent to SGPs compared to wild type (Fig. 4C). In *epi-1(RNAi)* larva, nidogen/NID-1^mNG^ also localized to the gonadal BM at levels similar to control larva and showed discontinuities similar to those we observed in EMB-9^mNG^ (Fig. 4D–F). Finally, we observed that in wild-type newly hatched, starved L1 larva, perlecan/UNC-52^mNG^ surrounded the gonad primordium and was also present at low levels in the extracellular space between the PGCs (Fig. 4G, arrowheads). In *epi-1(RNAi)* larvae, UNC-52^mNG^ still localized robustly to the gonadal BM at comparable levels to wild-type larvae (Fig. 4H,I). We conclude that *epi-1* RNAi does not prevent accumulation of type IV collagen, nidogen or perlecan and that these components could engage integrin receptors when laminin is knocked down.

**Figure 4.**
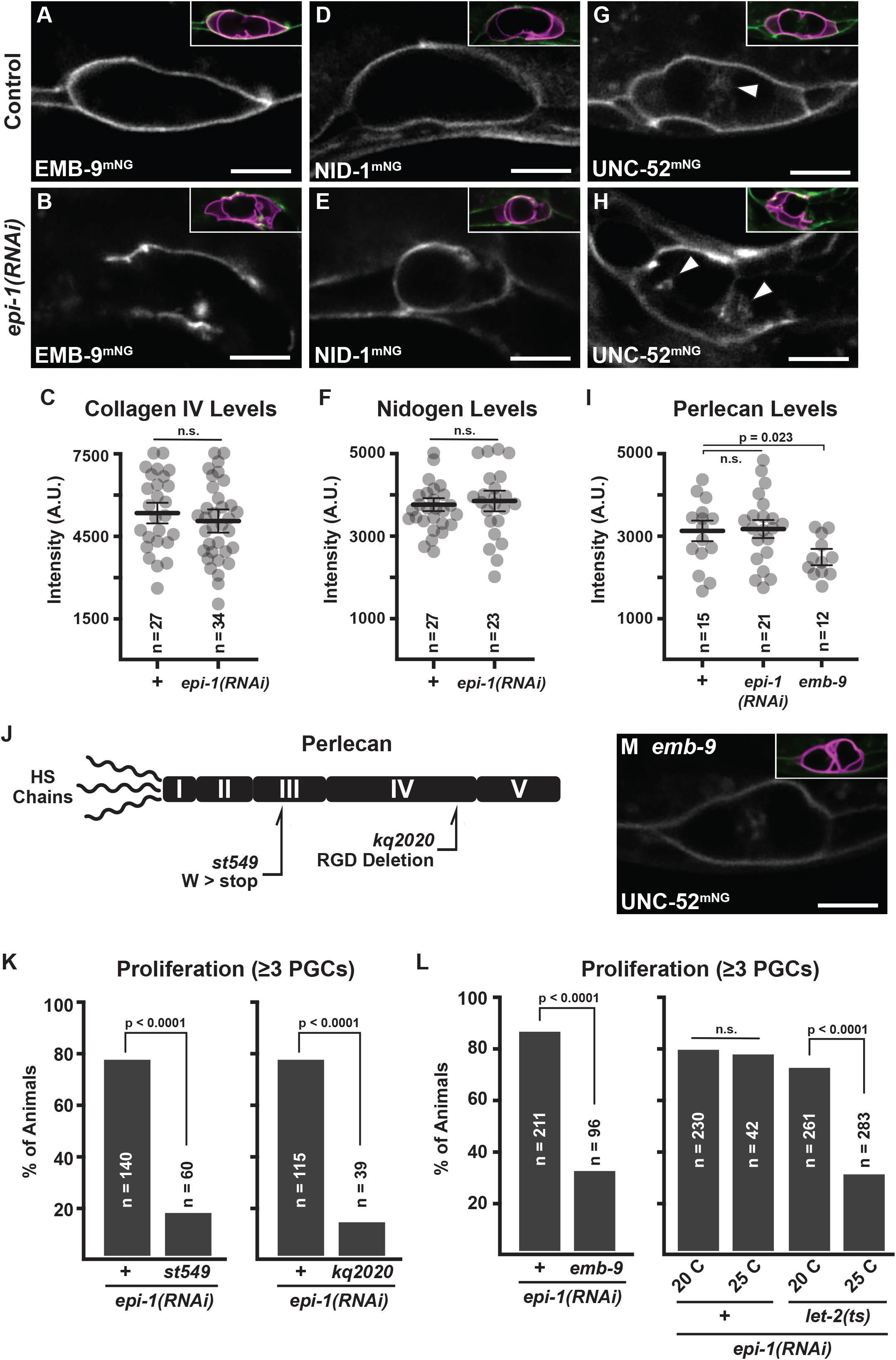
Perlecan is required for PGC proliferation when laminin is depleted. **A – B**. Collagen IV 1/EMB-9^mNG^ accumulation around L1 gonad primordia in control (A) and *epi-1(RNAi)* (B) animals. **C**. Quantification of mNG fluorescence in BM adjacent to SGP membranes from A and B. **D – E**. Nidogen/NID-1^mNG^ accumulation around L1 gonad primordia in control (D) and *epi-1(RNAi)* (E) animals. **F**. Quantification of mNG fluorescence in BM adjacent to SGP membranes from D and E. **G – H**. Perlecan/UNC- 52^mNG^ accumulation around L1 gonad primordia in control (G) and *epi-1(RNAi)* (H) animals. **I**. Quantification of mNG fluorescence in BM adjacent to SGP membranes from G, H and M. A – H, Insets show SGP membranes (magenta) with the indicated BM component (green). **J**. diagram showing perlecan domain structure, N-terminal heparan sulfate modifications and locations of the null (*st549*) and RGD deletion (*kq2020*) mutations tested. **K – L**. Quantification of the fraction of 2 day starved L1 larvae with proliferating PGCs for the genotypes indicated. **M**. Perlecan/UNC-52^mNG^ accumulation around L1 gonad primordia in *emb-9* mutants. Scale bars are 5μm and specific p-values are listed (n.s., not significantly different, p > 0.05). Fisher’s exact test was used to compare the faction of worms with proliferating PGCs in K – L. All other quantitative comparisons used a 2-tailed t-test.

Both perlecan and collagen IV are ligands for RGD integrin receptors. Perlecan/UNC-52 is a ligand for PAT-2/PAT-3 integrin receptors in *C. elegans* body-wall muscle cells, and *unc-52* mutants, like *pat-2* and *pat-3* mutants, arrest at the 2-fold stage (Williams and Waterston, 1994, Rogalski et al., 1993, Guo et al., 1991, Gettner et al., 1995). Likewise, collagen IV is recruited to pharyngeal BM by PAT-2/PAT-3 integrins (Jayadev, JCB, 2019) (Jayadev et al., 2019). We hypothesized that either perlecan or collagen IV would be required for PGC proliferation when laminin is depleted. To test whether perlecan is required for PGC proliferation in laminin-depleted embryos, we examined *unc-52*(*st549)* mutants; *unc-52(st549)* contains a premature stop codon predicted to severely truncate the UNC-52 protein (Fig. 4J) (Williams and Waterston, 1994). Compared to control *epi-1(RNAi)* worms, PGC proliferation was suppressed in starved *unc-52(st549) epi-1(RNAi)* mutants (Fig. 4K) to nearly the same extent as *pat-3 epi-1(RNAi)* mutants (see Fig. 2F). Next, we tested if the collagen IV matrix is required for PGC proliferation by examining loss-of-function mutations of both of the collagen IV subunits: an engineered putative null allele of *emb-9* (*xn201*; Fig. S3C) and the *let- 2(b246)* temperature-sensitive lethal allele (Wood et al., 1980). Starved *emb-9 epi- 1(RNAi)* larvae had significantly fewer PGCs compared to *epi-1(RNAi)* alone (Fig. 4L). Likewise, *let-2(ts) epi-1(RNAi)* larva had significantly fewer PGCs when raised at the restrictive temperature (25°C) compared to larva raised at a permissive temperature (20°C) (Fig. 4L). However, in contrast to removing integrin (*pat-2* or *pat-3*) or perlecan (*unc-52*), loss of collagen IV only partially suppressed PGC proliferation. Together, these results indicate that perlecan plays an essential role in promoting PGC exit from quiescence when laminin is depleted, whereas collagen IV contributes to PGC quiescence but plays a modifying role.

Collagen IV can bind to perlecan, leading us to speculate that the weaker suppression phenotype of mutations disrupting collagen IV (*emb-9* and *let-2*) compared to those in perlecan (*unc-52*) could result from reduced levels of perlecan in the gonadal BM of collagen mutants. To test this hypothesis, we examined UNC-52^mNG^ levels in the gonadal BM of *emb-9* larvae (Fig. 4I,M). In contrast to control and *epi-1(RNAi)* larvae, which contained similar levels of UNC-52^mNG^ in the gonadal BM, levels of UNC-52^mNG^ in *emb-9* mutant larvae were moderately reduced. This finding suggests that collagen could contribute to PGC quiescence by promoting perlecan enrichment in the gonadal BM, although we cannot exclude a signaling role for collagen IV.

Since PAT-2/PAT-3 integrins are thought to interact with RGD peptides in target ligands, we tested whether a putative integrin-binding RGD sequence within perlecan/UNC-52 is important for regulating PGC quiescence. Targeted deletion of the second of two RGD sequences, within domain IV of UNC-52 *[unc-52(kq2020)]*; Fig. 4J) causes embryonic arrest at the two-fold stage (Qiu et al., 2022). This phenotype is shared with *unc-52* and *pat-3* putative null mutants, suggesting that the domain IV RGD functions as an essential integrin ligand. Deleting the domain IV RGD in domain IV suppressed PGC proliferation in *epi-1(RNAi)* animals to a similar extent as did the *unc- 52(st549)* null mutation (Fig. 4K). Because previous antibody staining experiments indicate that UNC-52 protein is still produced and localizes in *unc-52(kq2020)* mutants (Qiu et al., 2022), these experiments suggest that the domain IV RGD of UNC-52 may function as an integrin-binding motif required to trigger PGC proliferation when laminin is depleted.

## Discussion

Many studies have demonstrated that basement membrane signals are essential for proper regulation of stem cell survival, self-renewal and differentiation (Suh and Han, 2011, O’Reilly et al., 2008, Fujiwara et al., 2011). However, BMs are dynamic, heterogenous structures whose composition varies not just across time and space but between healthy and diseased states (Tsutsui et al., 2021, Keeley et al., 2020, Kai et al., 2019, Ge et al., 2020). Comparatively little is known about how cells respond to changes in the composition of adjacent BMs, nor about how such changes are regulated during development. The data we have presented show that depleting laminin changes the composition, and ultimately the function, of the gonadal BM. Relying on endogenously tagged BM components and engineered mutant alleles, we find that knocking down laminin α/*epi-1* does not prevent assembly of collagen IV, nidogen or perlecan to the gonadal BM. Perlecan and collagen IV are in fact *required* for PGCs to exit quiescence when laminin α/*epi-1* is knocked down. Under this model, niche cells act as intermediaries that convey information from the BM to the PGCs (Fig. 5A). Laminin in the gonadal BM normally prevents perlecan from engaging integrin receptors in SGP niche cells (Fig. 5B). When this inhibition is relieved, perlecan activates integrin receptors on SGP niche cells (Fig. 5C). This in turn activates Notch signaling in PGCs, inducing them to exit from quiescence and begin to divide.

**Figure 5.**
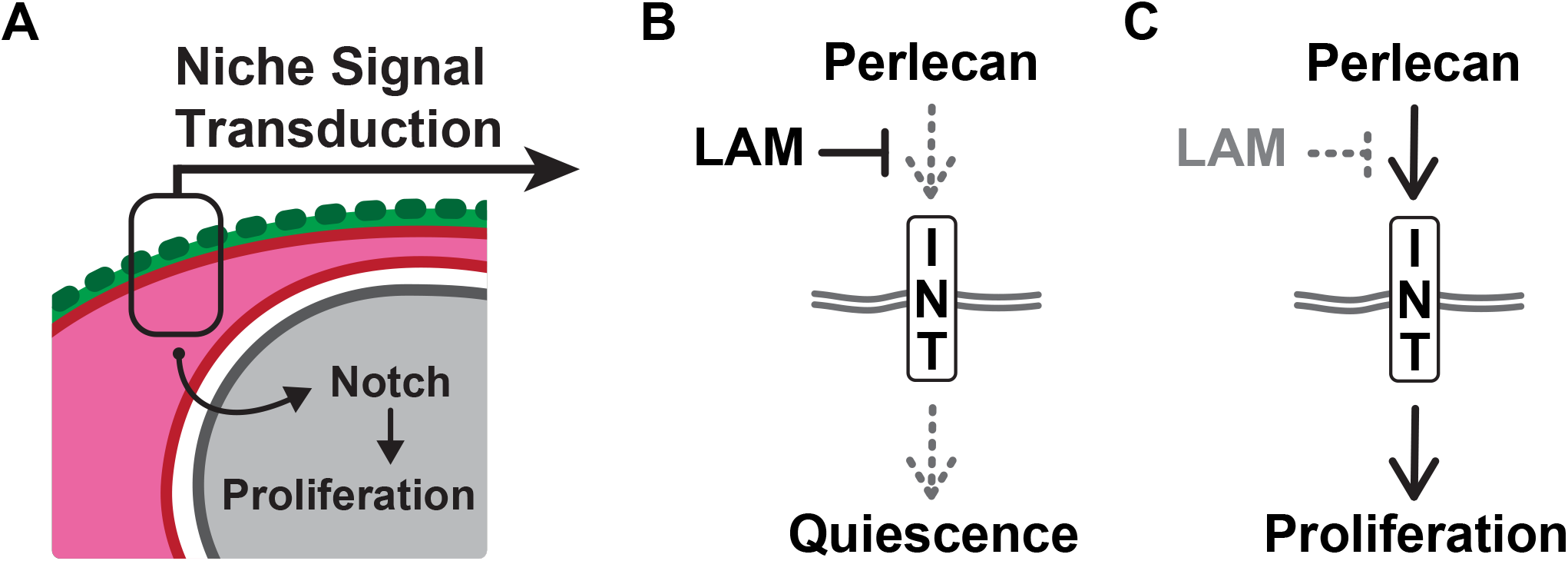
SGPs regulate PGC quiescence in response to changes in basement membrane composition. **A**. SGP niche cells (magenta) function as intermediaries that convey information from the BM (green) to the PGCs (gray). **B**. Model proposing that laminin (LAM) blocks activation of integrin (INT) by perlecan, maintaining PGC quiescence. **C**. Model proposing that in the absence of laminin, engagement of integrin through perlecan causes PGCs to exit quiescence and begin to proliferate.

There are several possible ways that altering gonadal BM composition could affect niche function. One way this may occur is if changes in the mechanical properties of the BM facilitate perlecan interactions with integrin. In both developmental and cancer models, increasing the amount of collagen IV relative to laminin stiffens BM and can drive cell proliferation (Kai et al., 2019). Alternatively, the BM could be so compromised in *epi- 1(RNAi)* larvae that normal mechanical cues are lost. Similar mechanisms have been reported in mammalian cells, where integrin αV signaling is need to maintain the quiescence of hematopoietic stem cells in the bone marrow and spleen, and integrin-mediated adhesion to osteopontin supports quiescence of disseminated acute lymphoblastic leukemia cells (Mehatre et al., 2021, Khurana et al., 2016, Horton et al., 2020, Boyerinas et al., 2013). A distinct possibility is that laminin knock down liberates perlecan from the BM. In the *Drosophila* ovary, perlecan functions both within the BM and the interstitial matrix to maintain the germline stem cell pool (Diaz-Torres et al., 2021). We observed interstitial perlecan within the gonad in laminin-depleted worms, raising the possibility that BM-free perlecan could contribute to proliferation. Finally, altering niche BM composition could change the way that perlecan interacts with one of the numerous growth factors that it binds. Often this cooperation controls stem cell proliferation. For example, perlecan cooperates with FGF to activate signaling via the Ras/ERK or PI3K/AKT pathways leading to increased proliferation of mammalian chondrocytes and neural stem cells (Vincent et al., 2007, Smith et al., 2007b, Smith et al., 2007a, Patel et al., 2007, Kerever et al., 2007, Kerever et al., 2014, GHISELLI et al., 2001). Changes to BM composition therefore could impact the availability of growth factors in the niche, leading to PGC proliferation. Since perlecan interacts with Wnt and TGFβ signals during cell migration as the gonad grows in larva (Merz et al., 2003), this mechanism could explain perlecan’s ability to trigger PGC proliferation when laminin is knocked down.

Studies in other systems have indicated that integrin activity is used to tune Notch signaling to the local ECM environment. For example, integrin promotes Notch signaling in order to control endothelial-to-mesenchymal transition following blood vessel damage (Wang et al., 2018b), differentiation of hematopoietic stem cells (Hadland et al., 2022), self-renewal of mouse embryonic stem cells (Suh and Han, 2011) and proliferation of neural stem cells (Shen et al., 2004, Rosa et al., 2016, Androutsellis-Theotokis et al., 2006). Conversely, activation of integrin by extra-cellular MAGP2 *negatively* regulates Notch in endothelial cells (Deford et al., 2016). In the worm gonad primordium, integrin also appears to coordinate Notch signaling with the local ECM environment. However, modulation of Notch signaling in the gonadal niche differs from these examples because integrin signaling in SGPs must activate Notch signaling in PGCs non-autonomously. How this occurs will be an important area of future study. For example, integrin signaling could promote the expression of a Notch ligand in SGPs, or alter its activity or localization. Compared to regulation of Notch receptors, less is known about how integrin signaling modulates the activity of Delta-like Notch ligands. One example where this has been investigated is the branching morphogenesis of new blood vessels. Tip cells signal to surrounding endothelial cells via Delta-Notch, and Notch activation favors vessel elongation and inhibits new vessel branches from forming. Laminin in surrounding BMs contributes to Notch activation by inducing expression of Delta/DLL4 (in tip cells) in an integrin β1-dependent pathway (Stenzel et al., 2011, Estrach et al., 2011). Our results identify PAT-2/PAT-3 integrin heterodimers as receptors needed in niche cells in order to tune notch signaling with BM composition. However, the signal transduction mechanism(s) downstream of integrin in the gonad have yet not been identified.

Uncovering these mechanisms will also illustrate how BM cues are integrated with nutritional signals to control stem cell quiescence and the onset of proliferation. Independent nutritional signaling through PI3 Kinase/AGE-1, PTEN/DAF-18 and AKT-1 also controls PGC escape from quiescence in response to food (Hubbard and Schedl, 2019, Fukuyama et al., 2006, Fry et al., 2021). Since disruption of gonadal BM bypasses nutritional regulation and operates independently of *daf-18* (McIntyre and Nance, 2020), niche cells may coordinate these two mechanisms in order to determine when PGCs exit quiescence. Coordination could happen at the level of signal transduction, similar to many cancers where integrin cooperates with receptor tyrosine kinases to drive proliferation though PI3K and AKT (Cooper and Giancotti, 2019). BM and nutritional signals could also act in parallel to regulate a common target ligand. In fly neuroblasts, nutritional signals (via PI3K and AKT) regulate stem cell exit from quiescence when larva begin to feed (Chell and Brand, 2010, Otsuki and Brand, 2018). However, in mushroom body neuroblasts, Pax-6/eyeless expression decouples stem cell proliferation from nutrition, similar to loss of laminin in the worm gonad (Sipe and Siegrist, 2017). Perhaps integrin signaling activates a transcription factor like VAB-3/Pax-6, which does function in *C. elegans* distal tip migration (Meighan and Schwarzbauer, 2007), to induce Notch ligand expression.

Finally, a significant implication of our findings is that niche cells interpret environmental cues, including BM composition, in order to modulate stem cell behavior. Laser ablation studies show that PGCs do not proliferate without somatic niche cells (McIntyre and Nance, 2020, Kimble and White, 1981). In addition, our findings using tissue-specific degradation demonstrated that integrin is needed in SGP niche cells to trigger PGC proliferation when laminin α/*epi-1* is depleted. Therefore, in contrast to many other types of stem cells, which either respond directly to nutritional signals (such as *Drosophila* neuroblasts) (Chell and Brand, 2010) or physically interact with BM (such as *Drosophila* intestinal stem cells) (You et al., 2014), *C. elegans* niche cells sense the extracellular environment and modulate signaling to the PGCs accordingly. Because of its ability to interact with both integrin and various growth factor signaling pathways, perlecan may function as a critical node in this process. Its necessity, along with integrin, demonstrates the importance of changes to BM composition in determining when quiescent stem cells are permitted to proliferate.

## Methods

### KEY RESOURCES TABLE

#### Bacterial and Virus Strains

| Reagent or Resource | Source | Identifier |
| --- | --- | --- |
| OP50, *E. coli* | *Caenorhabditis* Genetics Center | OP50 |
| RNAi feeding, *E. coli* | *Caenorhabditis* Genetics Center | HT115 |

#### Chemicals, Peptides, and Recombinant Proteins

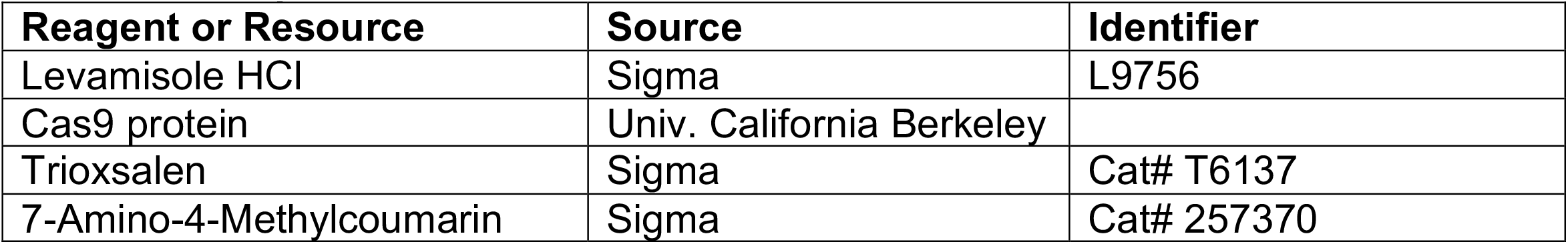

#### Experimental Models: Organisms/Strains

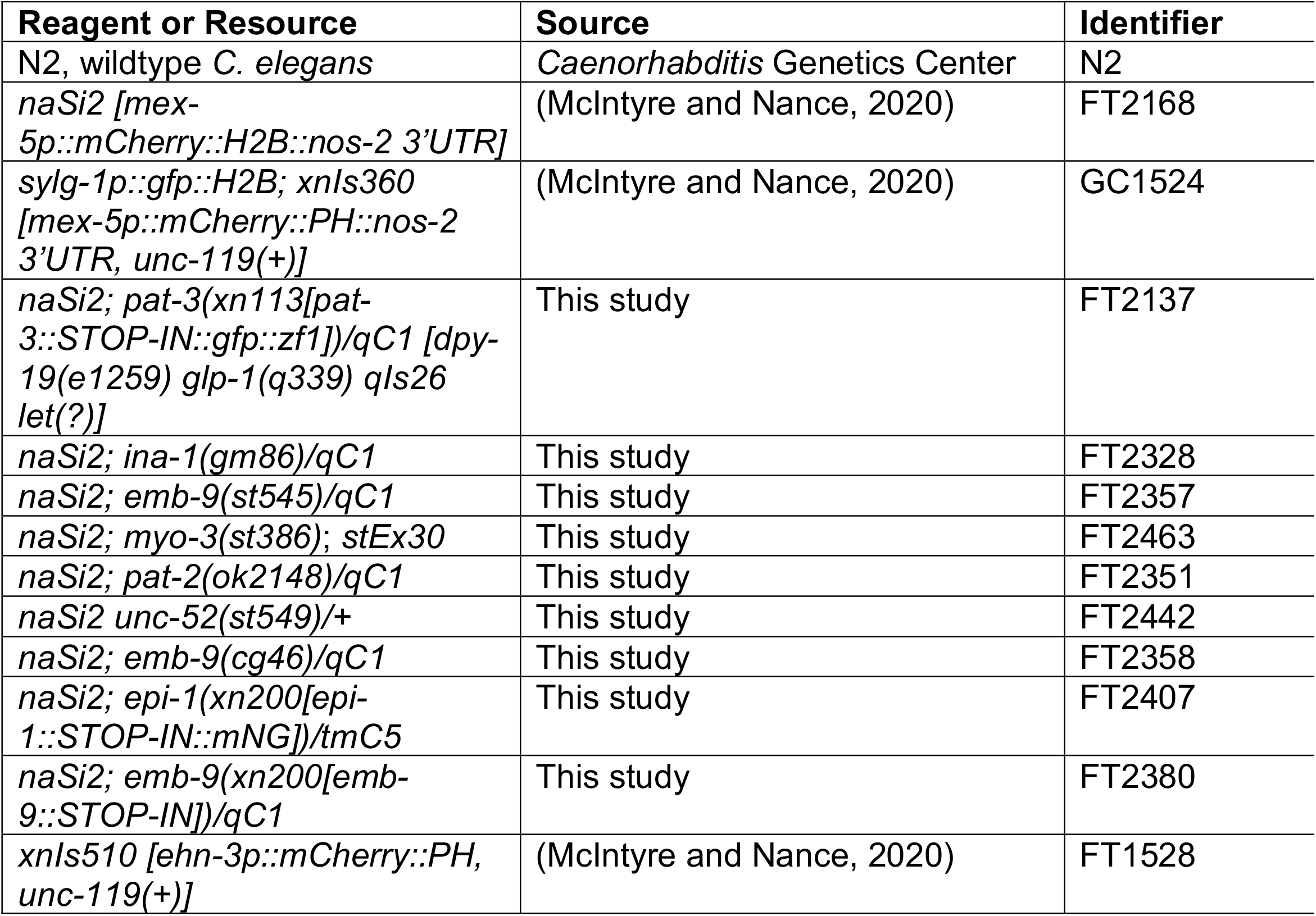
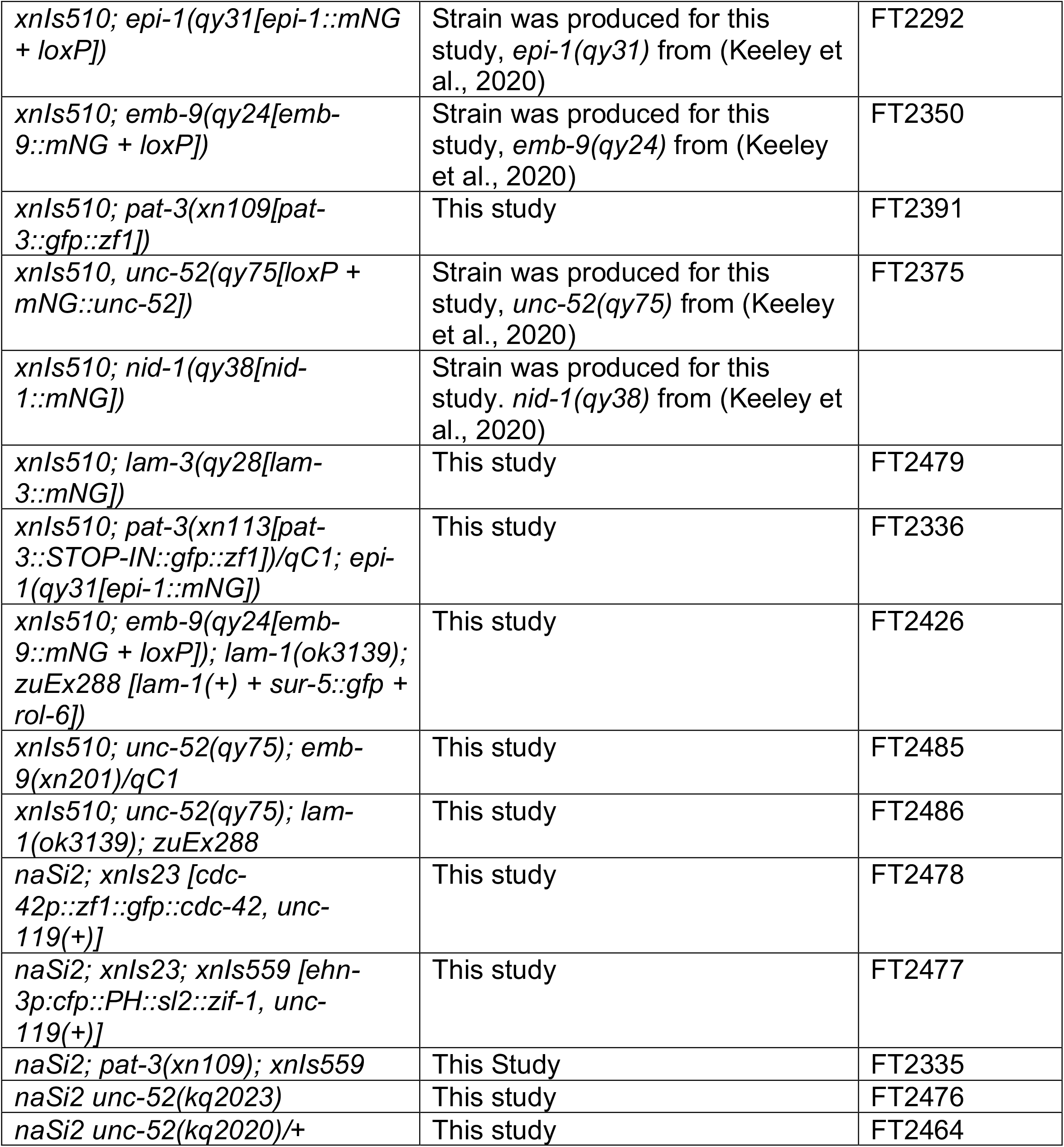

#### Oligonucleotides

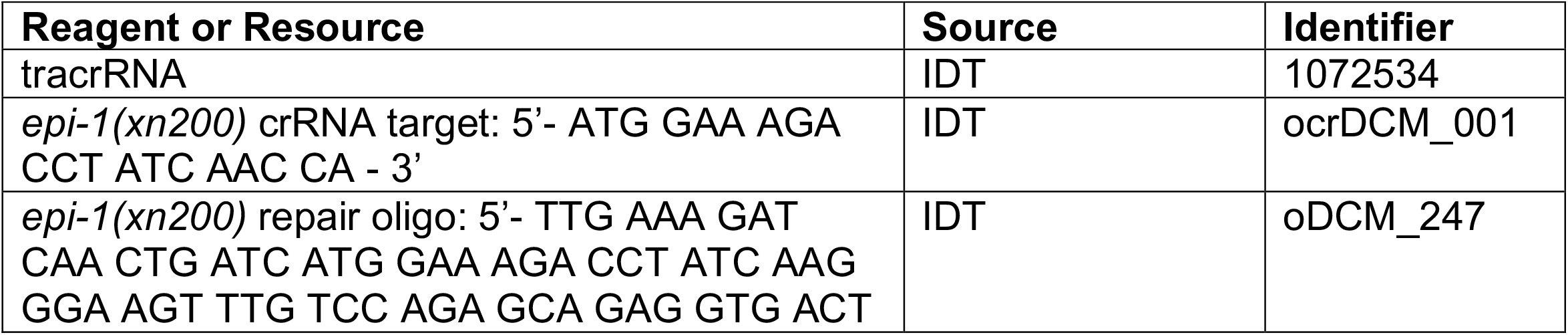
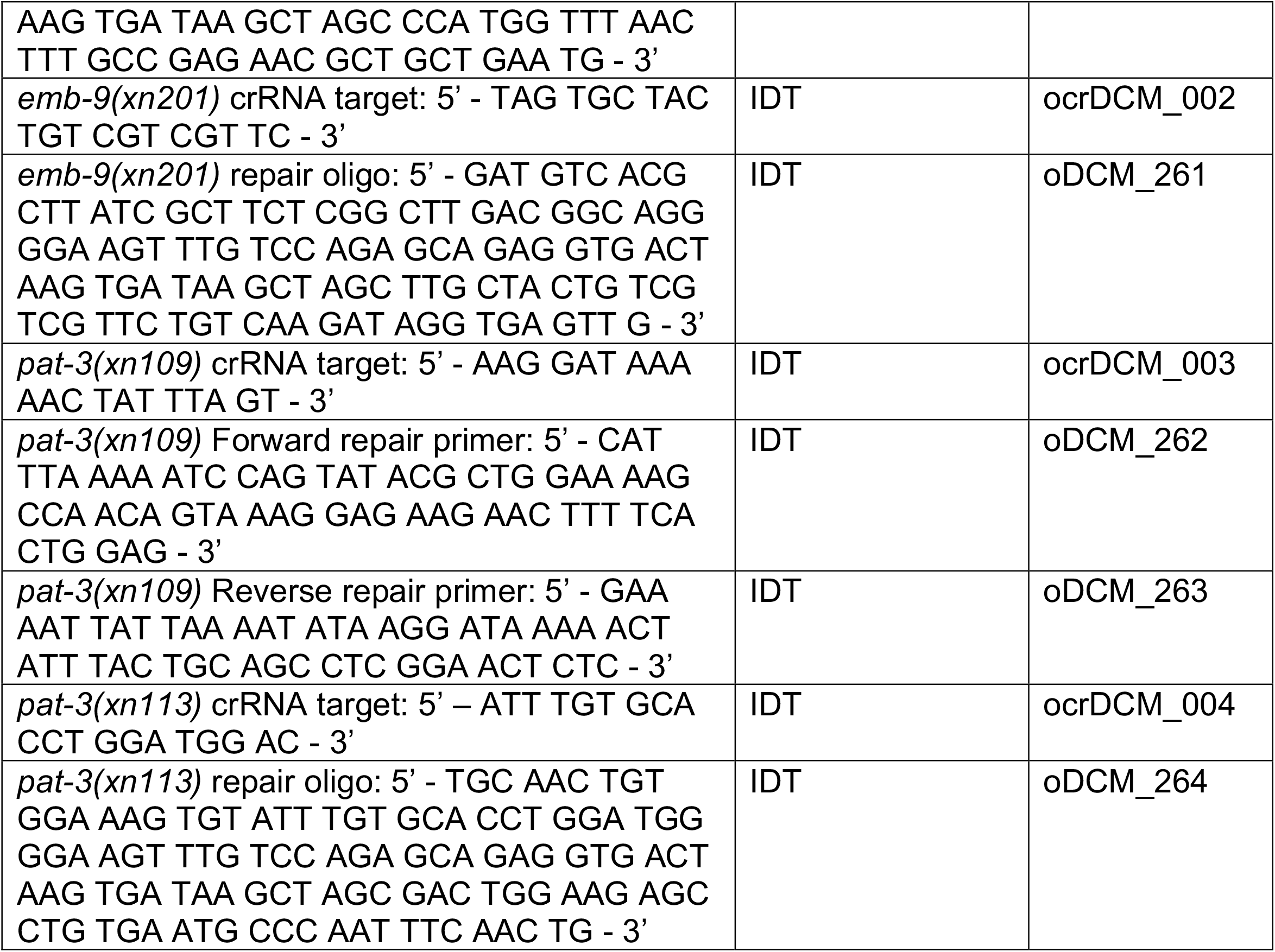

#### Recombinant DNA

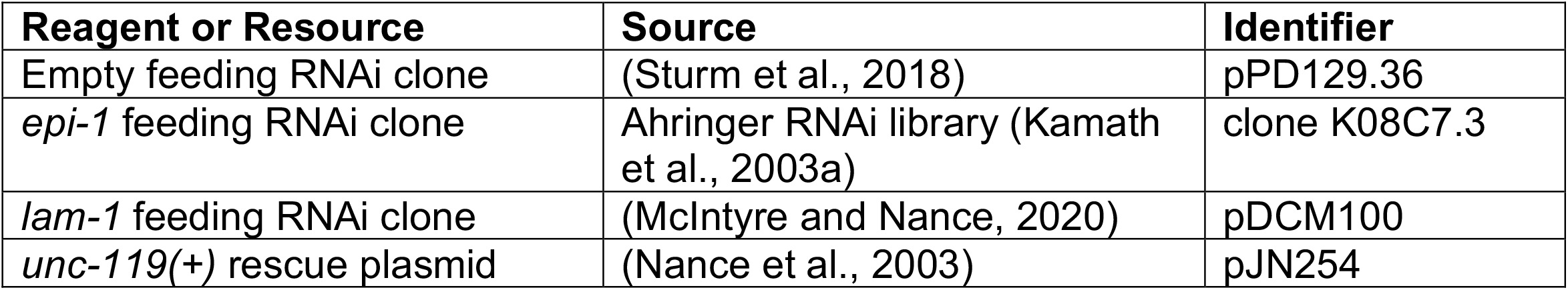

| Reagent or Resource | Source | Identifier |
| --- | --- | --- |
| Empty feeding RNAi clone | (Sturm et al., 2018) | pPD129.36 |
| *epi-1* feeding RNAi clone | Ahringer RNAi library (Kamath et al., 2003a) | clone K08C7.3 |
| *lam-1* feeding RNAi clone | (McIntyre and Nance, 2020) | pDCM100 |
| *unc-119(+)* rescue plasmid | (Nance et al., 2003) | pJN254 |

#### Software and Algorithms

| Reagent or Resource | Source | Identifier |
| --- | --- | --- |
| FIJI v2.0 | FIJI | https://fiji.sc/ |
| Zeiss Zen software v2.0 | Carl Zeiss | https://www.zeiss.com/microscopy/en/products/software/zeiss-zen.html |
| Nikon Elements | Nikon | https://www.microscope.healthcare.nikon.com/products/software/nis-elements |
| LAS X | Leica | https://www.leica-microsystems.com/products/microscope-software/p/leica-las-x-ls/ |
| Adobe Photoshop CC v19.0 | Adobe Systems | https://www.adobe.com/products/photoshop.html |
| Adobe Illustrator CC v19.0 | Adobe Systems | https://www.adobe.com/products/illustrator.html |

### EXPERIMENTAL MODEL SYSTEM

#### *C. elegans* Growth and Culture

Worms were cultured at 23°C under standard experimental conditions (Brenner, 1974) unless otherwise described. Strain N2 (Bristol) was used as the wild type. Genotypes of strains utilized in this study are listed in the Key Resources Table.

### TRANSGENESIS AND GENOME EDITING

#### Transgenes

*ehn-3p::cfp::PH::sl2::zif-1::tbb-2 3’ UTR* (pJN640) was created by multi-fragment Gibson assembly using vector backbone pCFJ151 (Frokjaer-Jensen et al., 2008), which contains the *unc-119(+)* co-transformation marker. Plasmid DNA was injected into *unc- 119(ed3)* worms to obtain an extrachromosomal array, which was integrated using Trioxsalen and UV irradiation as described (Armenti et al., 2014). The resulting multicopy transgene insertion, *xnIs559*, was outcrossed twice before use.

#### CRISPR Genome Engineering

CRISPR/Cas9-mediated genome editing was used to engineer mutations into the genome, as described (Paix et al., 2017). Purified Cas9 protein was pre-incubated with crRNA and tracrRNA, mixed with ssDNA oligonucleotide or dsDNA PCR repair templates containing >35bp homology arms, and injected into worms together with *dpy-10* co-CRISPR reagents. F1 Roller worms were singled and genotyped by PCR to identify edited worms. crRNA, repair template sequences and genotyping primers are listed in the Key Resources Table.

##### epi-1(xn200[epi-1::STOP-IN::mNG+loxP])

To make an *epi-1* null allele, the STOP-IN cassette was inserted into second exon of the previously described *epi-1(qy31[epi-1::mNG+loxP])* allele (Keeley et al., 2020). *epi- 1(qy31)* was cut with crRNA ocrDCM_001 and repaired with an ssDNA oligo (oDCM_247) that contained the STOP-IN cassette. *epi-1(xn200)* was maintained over the *tmC5* balancer (Dejima et al., 2018). See Supplemental Figure 1.

##### pat-3(xn109[pat-3::gfp::zf1])

To create the repair template, *gfp::zf1* lacking the initiator methionine was amplified by PCR from a plasmid template using primers oDCM_262 and oDCM_263, which contained homology arms to the *pat-3* 3’ end. *pat-3* was cut with crRNA ocrDCM_003 and repaired with the dsDNA PCR repair template, resulting in *gfp::zf1* insertion immediately after the final *pat-3* codon without a linker sequence. Insertion *pat- 3(xn109)* was verified by PCR, sequencing, and GFP expression. *pat-3(xn109)* animals were viable (n = 94/99 embryos developed to adult). See Supplemental Figure 2.

##### pat-3(xn113[pat-3::STOP-IN::gfp::zf1])

To create a putative *pat-3* null allele, we used the “STOP-IN” method (Wang et al., 2018a) to introduce a stop codon (in any reading frame) into the sixth exon, upstream of sequences encoding the transmembrane domain, in *pat-3(xn109[pat-3::gfp::zf1]). pat- 3(xn109)* was cut with crRNA ocrDCM_004 and repaired with ssDNA oligo (oDCM264), which contained the STOP-IN cassette. *pat-3(xn113)* was maintained over the *qC1* balancer. See Supplemental Figure 2.

##### emb-9(xn201[emb-9::STOP-IN::mNG+loxP])

To make an *emb-9* null allele, the STOP-IN cassette was inserted into the first exon of the previously described *emb-9(qy24[emb-9::mNG])* allele (Keeley et al., 2020). *emb-9(qy24)* was cut with crRNA ocrDCM_002 and repaired with an ssDNA oligo (oDCM_261) that contained the STOP-IN cassette. *emb-9(xn201)* was maintained over the *qC1* balancer. See supplemental Figure 3.

#### RNAi

*E. coli* bacterial strain HT115 expressing dsRNA from vector pPD129.36 or its derivatives was fed to L4 *C. elegans* larvae. Plates contained 0.2% ß-lactose to induce dsRNA production. L4 larval worms were fed for two days at 23°C before progeny were analyzed. To collect progeny, adult hermaphrodites were washed 2X, dissected in Egg Salts, and early-stage embryos were collected into fresh buffer. Embryos were then aged at 23°C for 2 days in Egg Salts (unless a different length of times is specifically indicated). Results shown include data from at least three independent trials. As a negative control, worms were fed empty pPD129.36 vector. The RNAi feeding plasmid targeting *epi-1* is Ahringer RNAi library clone K08C7.3 (Kamath et al., 2003b), and the RNAi feeding plasmid targeting *lam-1* was previously described (McIntyre and Nance, 2020). The effectiveness of *epi-1* RNAi was confirmed by comparison with the *epi-1(xn200)* putative null allele, as shown in Figure 1.

### MICROSCOPY

#### Live Imaging of Embryos and Larvae

For live imaging, embryos or larvae were mounted on 4% agarose pads in Egg Salts. Young larvae were immobilized with 1-2ul of 5mM levamisole applied to the coverslip before it was placed on the agarose pad. PGCs were counted using a Zeiss AxioImager A2 equipped with an AxioCam 503 mono CCD camera, Uniblitz shutter, and 63X 1.4NA or 40X 1.3NA objective. Embryos and larva were imaged live using a spinning disk confocal microscope (Nikon Eclipse Ti2, CSU-W1 spinning disk, 100X 1.35NA Silicone oil immersion objective, 488nm and 561nm lasers, matched Andor 888 Live EMCCD cameras). A Leica SP8 confocal microscope (63X 1.4NA oil immersion objective, Argon tunable and HeNe 594nm lasers, and HyD detectors) was used for quantifying CDC- 42::GFP::ZF1 degradation. Unless specified otherwise, the distance between slices in Z stacks was 500nm. Images were acquired using Zeiss Zen software, Nikon Elements or Leica Application Suite X before being processed in ImageJ and Adobe Photoshop.

#### Laser Ablation of Somatic Gonadal Precursor Cells

Ancestors of the SGPs (MSapp and MSppp) were ablated as described (McIntyre and Nance, 2020). Control unablated embryos were mounted on the same slides as experimental embryos and allowed to develop simultaneously. Rarely, laser ablation caused damage to nearby control unablated embryos. Such experiments were discarded and repeated.

### IMAGE QUANTIFICATION

#### Basement Membrane Protein Intensity Measurements

To compare the fluorescence intensity of BM components and receptors, confocal images of control and perturbed animals were first acquired using equal laser power and exposure times and comparable focal depths. The maximum pixel intensity (averaged over a line 10 pixels in width) was measured across the gonadal BM on the side of the somatic gonad facing the endoderm. Because background fluorescence (as measured in the PGC nucleus) was minor and very consistent across the data, background fluorescence was not subtracted prior to image intensity quantification.

#### Notch Reporter Nuclear Intensity Measurements

GFP::H2B expression from *sygl-1p::gfp::H2B* was quantified in the PGC nucleus by capturing images of control and SGP-ablated embryos with equal exposure settings using a Zeiss AxioImager A2 descibed above. Total pixel intensity was measured in a 4µm diameter circle centered on the PGC nucleus and background fluorescence was subtracted by measuring GFP intensity in a region of the same size outside of the PGC.

### STATISTICAL ANALYSIS

Statistical tests used are indicated in figure legends. For categorical data, such as the fraction of worms with more than two PGCs, p-values were calculated using Fisher’s Exact Test. Other datasets, such as fluorescence intensity measurements, were analyzed using a two-tailed Student’s t-test. For t-test comparisons between two or more samples, normality was assessed according to the Shapiro-Wilk normality test. If multiple comparisons were made within a dataset, p*-*values were adjusted using the Bonferroni correction. Datasets presented represent at least three independent experiments with sample size indicated in each figure. Damaged embryos were excluded from analysis.

## Acknowledgements

We thank David Sherwood and Myeongwoo Lee for sharing *C. elegans* strains with our lab. We thank members of the Nance lab as well as E. Jane Hubbard and Niels Ringstad labs for discussion of this project and comments on the manuscript. We also thank Michael Cammer and Yan Deng from the NYU Langone Microscopy Laboratory, which is partially supported by the Cancer Center Support Grant P30CA016087 at the Laura and Isaac Perlmutter Cancer Center, for their advice and discussions of live imaging. Some strains were provided by the *Caenorhabditis* Genetics Center, which is funded by NIH Office of Research Infrastructure Programs (P40OD010440).

## Data availability

All materials described in this manuscript are either commercially available or available from the authors upon request. Worm strains are available from the *Caenorhabditis* Genetics Center or by request from the authors. No datasets were generated that should be deposited in public databases. Raw data are available upon request.

---

**Supplemental Figure 1. Construction of a laminin /*epi-1* null mutant. A**. A stop cassette was inserted into the second exon of laminin /*epi-1(qy31[epi-1::mNG+LoxP])* to generate *epi-1(xn200[epi-1::STOP-IN::mNG+LoxP])*. **B – C**. EPI-1^mNG^ fluorescence (green), with SGP membranes (magenta), in *epi-1(qy31)* control (B) and *epi-1(xn200)* mutant (C) L1 larvae. **D**. Quantification of the fraction of control and *epi-1* larvae that survive to adult. **E – F**. DIC images of control and *epi-1(xn200)* newly hatched L1 larva. *epi-1* mutant larva are kinked and often have a prominent nodule in the head region (arrowhead). Scale bars are 10μm.

**Supplemental Figure 2. Construction of integrin ß/*pat-3* mutant alleles. A**. GFP-ZF1 was inserted at the 3’ end of *integrin ß/pat-3* to generate *pat-3(xn109[pat-3::GFP::ZF1])*. **B**. A stop cassette was inserted into the sixth exon of *pat-3(xn109)* to generate *pat- 3(xn113[pat-3::STOP-IN::GFP::ZF1])*. **C**. Quantification of CDC-42^GFP-ZF1^ degradation using the ZIF-1^SGP^ driver. CDC-42^GFP-ZF1^ was quantified along SGP-PGC contacts (between the white arrow heads, also enlarged in insets). **D**. Average number of PGCs for *epi-1(RNAi)* and *myo-3; epi-1(RNAi)* in starved L1 larvae; colored circles indicate the mean in three independent experiments, and the bar is the mean of means. Scale bars are 5μm and specific p-values are listed (n.s., not significantly different, p > 0.05). Quantitative comparisons used a 2-tailed t-test.

**Supplemental Figure 3. Construction of a collagen IV α1/*emb-9* null allele. A**. A stop cassette was inserted into the first exon of *emb-9(qy24[mNG+LoxP::emb-9])* to generate *emb-9(xn201[STOPIN::mNG+LoxP::emb-9])*.

