## Supplementary figures and images for "Niche cells regulate primordial germ cell quiescence in response to basement membrane signaling"

### Supplemental Figure 1

#### Supplemental Figure 1

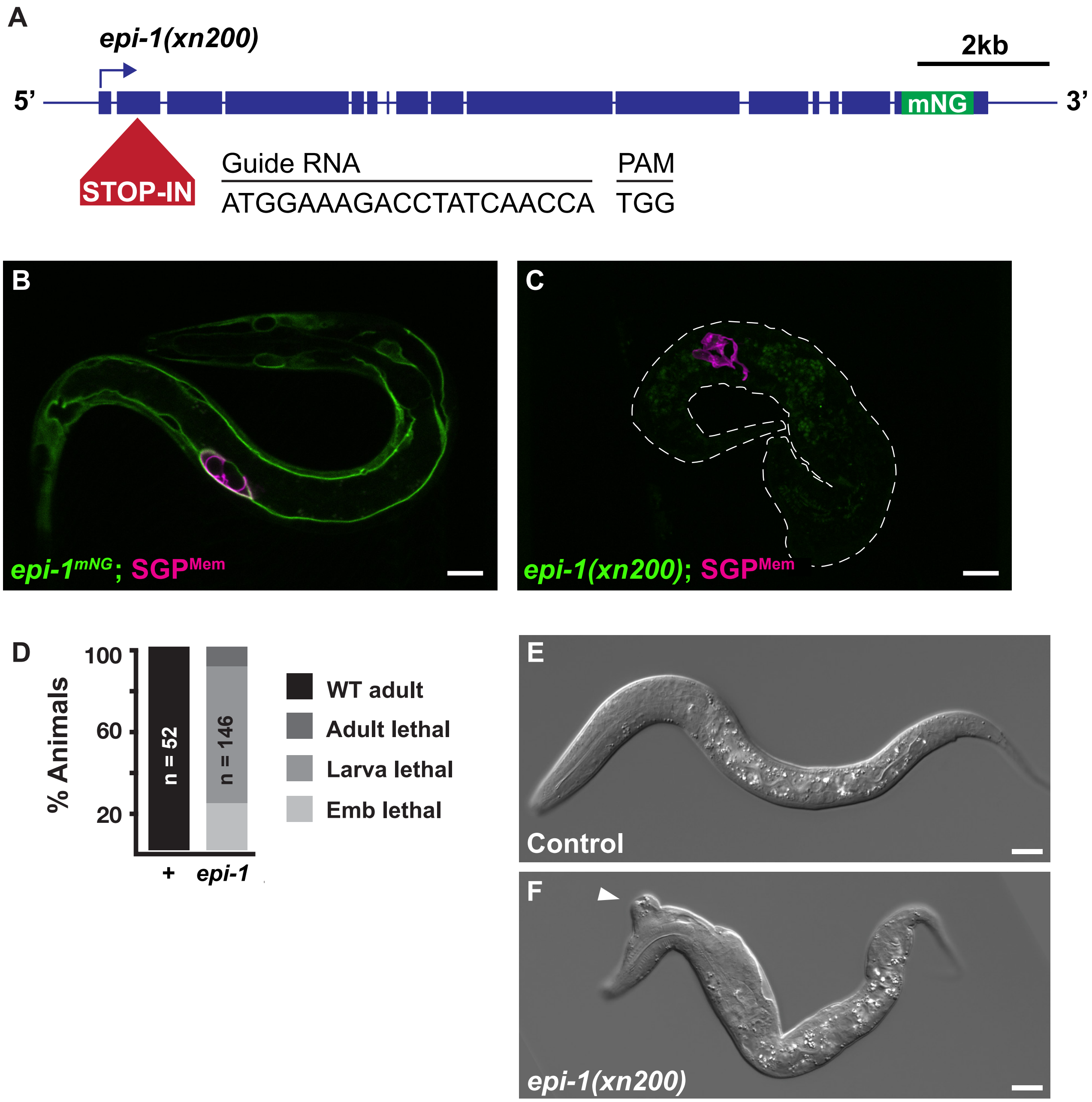

### Supplemental Figure 2

#### Supplemental Figure 2

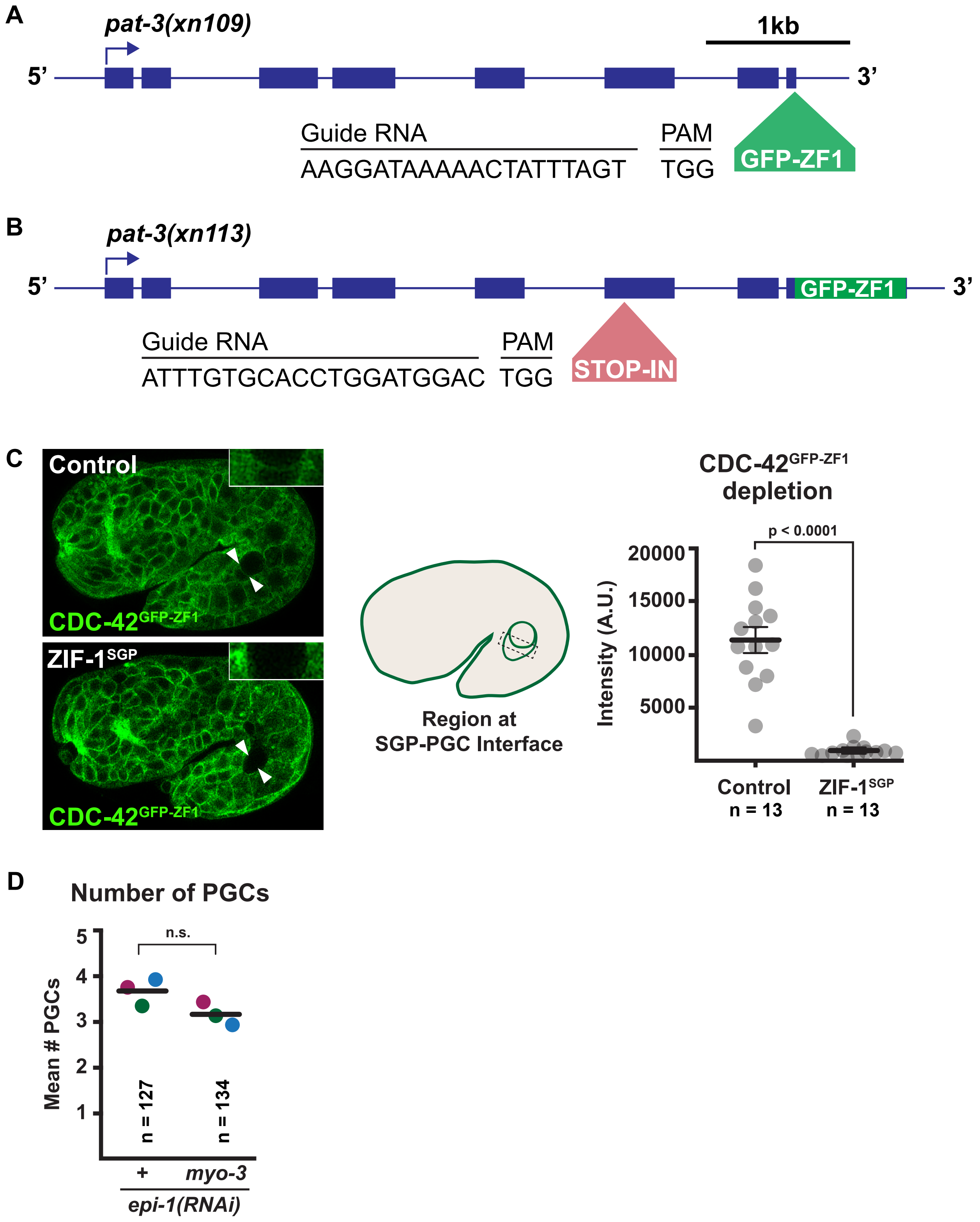

### Supplemental Figure 3

#### Supplemental Figure 3

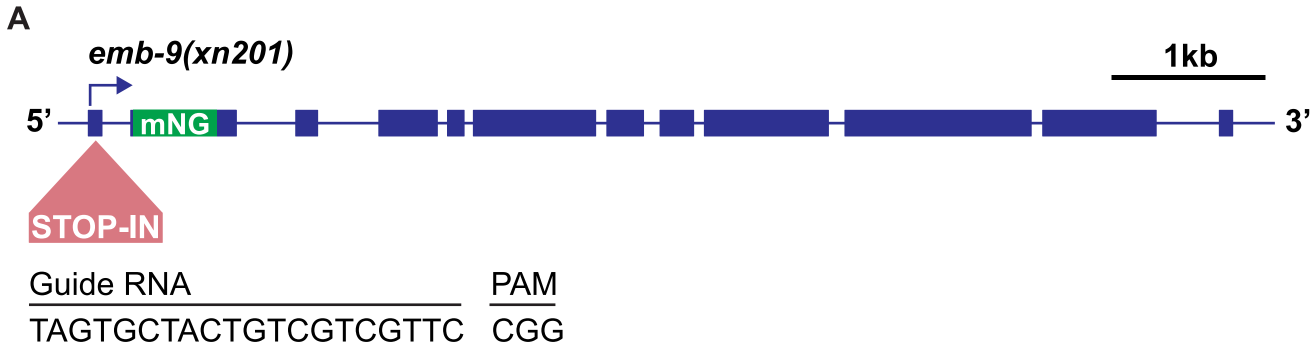
